# Improving Olfactory Receptor Structure Modeling via Hybrid Methods

**DOI:** 10.1101/2024.05.30.596580

**Authors:** Bhavika Berwal, Pinaki Saha, Ritesh Kumar

## Abstract

Understanding the structure of Olfactory Receptors (ORs) is pivotal in deciphering the molecular complexities of smell, a sense crucial for health, and survival, and holding immense therapeutic potential. However, the scarcity of detailed experimental data on ORs has hindered progress, demanding inventive approaches.

This study explores various structure prediction methods tailored to ORs based on their phylogenetic and structural characteristics, using OR51E2 as a reference. We employed a hybrid homology modeling approach, utilizing AlphaFold structures, yielding models with lower RMSD (1.019 ^°^A across pruned/significant pairs and 2.33 ^°^A over-all) and enhanced structural metrics compared to standalone AlphaFold (RMSD - 2.5 ^°^A) predictions. Our pipeline successfully replicated experimental findings for OR51E2 and was applied to homologous ORs: OR51E1, OR51D1, and OR51G2. Various tools were also used to predict potential binding sites for each receptor. Molecular dynamics simulations validated the stability of these OR models in a lipid bilayer environment, with biophysical analyses revealing that AlphaFold models exhibit relatively less ideal behavior compared to the Hybrid Model. Our study presents a targeted approach to investigate and generate optimum OR structures for further conformational analyses.

## Introduction

Olfaction, commonly known as the sense of smell, is an essential biological function that profoundly impacts the behavior and survival of various species, including humans. It plays a crucial role in detecting food, identifying environmental hazards, and navigating social interactions and reproductive behaviors. At the molecular level, olfaction is facilitated by the interaction of odorant molecules with the olfactory receptors (ORs). They are the largest family of G protein-coupled receptor (GPCR) family and are found within the Olfactory Sensory Neurons (OSNs) in the nasal epithelium.^1^ However, recent studies have revealed that olfactory receptors are not confined to the nasal cavity. These receptors are found in a variety of non-olfactory tissues and organs, where they perform a myriad of biological functions that extend beyond the conventional scope of olfaction.^2–4^ Their presence in extranasal tissues suggests functions beyond odor detection, including roles in fluid balance, wound healing, digestive processes, drug metabolism, respiratory control, and potentially behavior and mood regulation. This expanded understanding of ORs opens new research boundaries in molecular biology, highlighting their significance beyond traditional olfactory perception.^5,6^

Olfactory perception is so intricate that we have about 800 olfactory receptor proteins out of which about 400 are functional ORs, involved directly in the reception of odorants and perception of smell, thereby reflecting the complexity and diversity of our olfactory system.^7^ Understanding olfaction at the molecular level is significant for various reasons. Knowledge of olfactory receptors and their ligands holds immense potential in practical applications. The study of olfaction contributes to our broader understanding of GPCRs, which represent a major class of drug targets. GPCRs are also implicated in numerous physiological processes, and approximately 34% of all FDA-approved drugs target these receptors.^8,9^

Olfactory Receptors, like all GPCRs are are membrane proteins and consist of membranebound alpha helices, called Transmembrane helices. These proteins are often called Transmembrane proteins as well. ^10^ Membrane proteins, like GPCRs are known to have several activation states.^11^ Usually, protein structure prediction algorithms predict the inactive state of the receptor, which is the most stable conformation. However, understanding the active state of the receptor is crucial for drug discovery and understanding the mechanism of action of these receptors.

Our hybrid homology modeling approach, which combines the strengths of AlphaFold and traditional homology modeling, allows us to predict the active state of ORs with high accuracy. By leveraging the structural information from mammalian AlphaFold, specifically that of mouse OR structures and carefully selecting templates based on phylogenetic and structural characteristics, we can generate models that closely resemble the active state of the receptor. The importance of computational analysis in the study of GPCR structures cannot be overstated. Experimental techniques, such as X-ray crystallography and cryo-electron microscopy, have provided invaluable insights into the structure and function of GPCRs. However, these methods are time-consuming, expensive, and often require extensive optimization for each individual receptor.^12^ Moreover, capturing the active state of GPCRs experimentally is particularly challenging due to their inherent flexibility and the transient nature of their active conformations.^13^ Computational methods, such as homology modeling and molecular dynamics simulations, offer a complementary approach to experimental techniques. These methods allow researchers to predict the structure of GPCRs based on the available experimental data and to study their dynamics and interactions with ligands.^14,15^ However, the accuracy of these predictions relies heavily on the quality of the input data and the underlying assumptions of the computational methods.

Our hybrid homology modeling approach addresses these limitations by leveraging the strengths of both AlphaFold and traditional homology modeling. AlphaFold has revolutionized the field of protein structure prediction by using deep learning to generate highly accurate models.^16^ However, its performance on GPCRs, particularly in predicting active state conformations, has been less extensively validated.^17^ By carefully selecting templates based on phylogenetic and structural characteristics and incorporating mammalian AlphaFold structures, we can refine the computational predictions and generate models that more closely resemble the active state of the receptor.

This refined computational approach is particularly important for ORs, which are the largest class of GPCRs in mammals and play a crucial role in olfaction.^18,19^ Understanding the structure and function of ORs is essential for unraveling the molecular mechanisms of smell and for developing novel therapies for olfactory disorders. However, the experimental determination of OR structures has been hindered by their low expression levels, high instability, and difficulty in crystallization.^20,21^ Further advancements were made with the integration of bioinformatic techniques like Next-Gen Sequencing (NGS), allowing for a detailed analysis of OR sequences.^22^ These approaches involve the alignment and comparison of OR sequences across different species, aiding in the identification of conserved regions and critical amino acids responsible for receptor activation. These techniques have enabled us to de-orphanize ORs with higher capabilities. NGS being a High-throughput method, often requiring large computational resources can often be unviable for completely unrelated ORs, demanding an extensive pre-analysis of selected ORs often leading to in-accuracy.^23,24^ Another significant stride in this field has been the application of molecular dynamics simulations to study the dynamic behavior of GPCRs in a membrane medium that can also be applied to study ORs, this has been already done for OR51E2.^25^ These simulations provide a time-resolved view of the conformational changes in ORs, offering deeper insights into the mechanisms of receptor activation and signal transduction. Structural prediction tools like Alphafold2^16^ have revolutionized our ability to predict the three-dimensional structures of proteins, including GPCRs, with high accuracy.

Our study capitalizes on these technological advances to investigate the structure and function of a specific subset of human olfactory receptors. Ever since the structure extraction of the OR51E2 (PDB:8F76),^26^ there is a possibility to model other closely related receptors with an ever-known higher accuracy. The extent of computational analysis in overcoming these experimental challenges provides a framework for predicting the active state structure of ORs. By combining advanced structure prediction methods with phylogenetic and structural information, we can generate high-quality models that can guide future experimental studies and accelerate the discovery of novel odorants and potential therapeutic targets.

## Materials and Methods

BLAST was performed on OR51E2(PDB:8F76) to find the OR sequences with the highest sequence similarity. Templates with sequence similarity greater than 30% are usually selected to perform homology modeling,^27^ we thus selected the top three most similar receptors to OR51E2, shown in Fig1, namely OR51E1, OR51D1, and OR51G2. Our obtained sequences are well beyond this threshold and this makes ideal templates.

**Figure 1:**
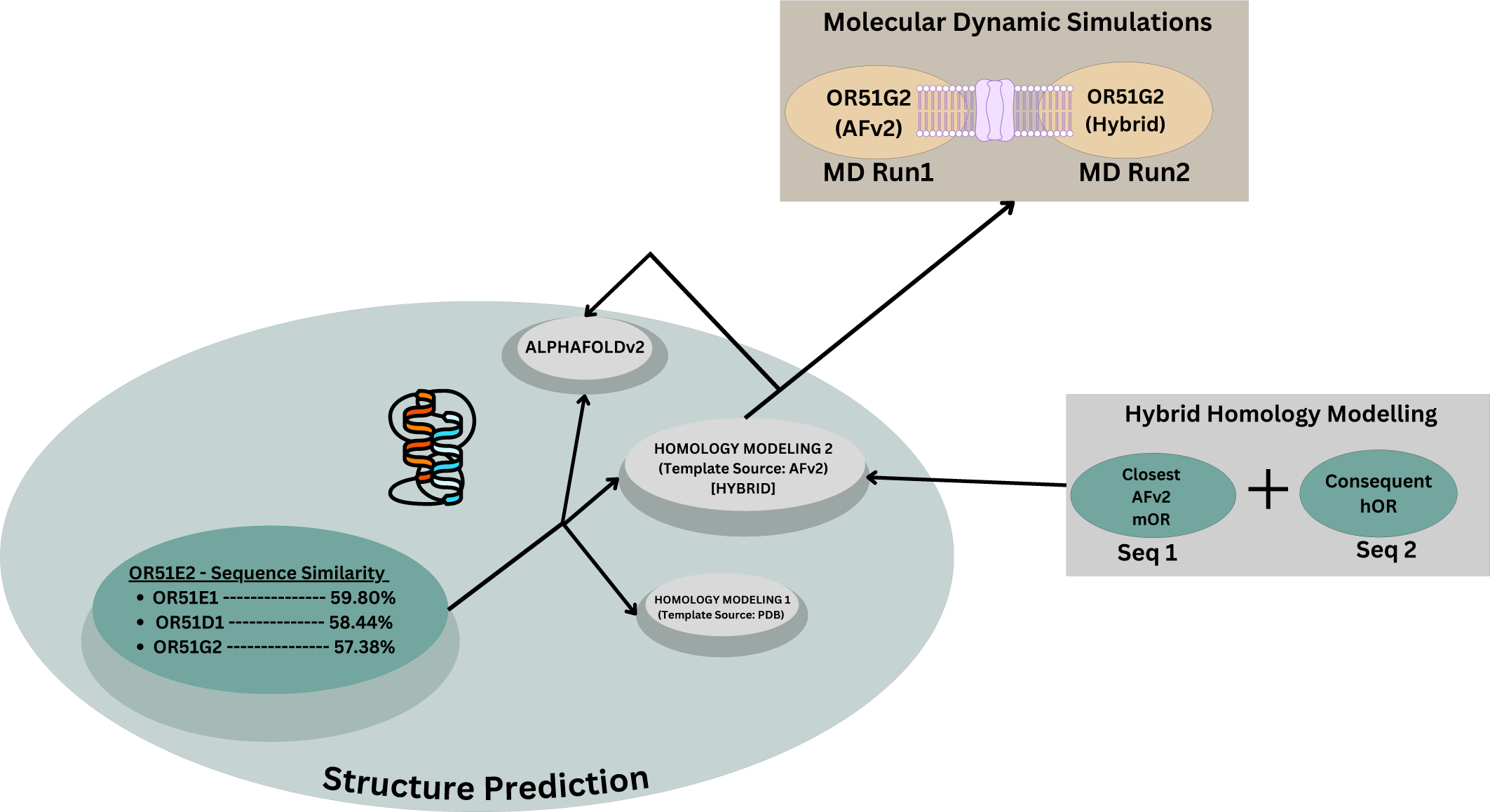
Flow-chart representing the methodology pipeline and Molecular Dynamics Simulations.

### Structure Prediction

To predict the structures of selected Olfactory Receptors (ORs), we employed three distinct methodologies. Initially, we utilized AlphaFold v2 to directly obtain the Protein Data Bank (PDB) structures from the EBI-AlphaFold database to assess the structure quality of AlphaFold-generated models under default settings. Traditional Homology Modeling was also applied via SWISS-MODEL, using OR51E2 as the template for modeling our selected ORs (OR51E1, OR51D1, and OR51G2) as this PDB protein shows the highest sequence similarity to these receptors.^28^ However, for modelling OR51E2 itself, we employed the human Cholecystokinin A receptor (CCKAR)-Gi complex (PDB: 7EZH) as a template, having the sequence identity of approximately 18% with OR51E2, falling below the typically required threshold for homology modeling *i.e.* 30%. This selection was based on the sequence with the closest structural resemblance to OR51E2, highlighting the inherent difficulty due to the scarce structural homology observed within GPCRs. Lastly, we explored a third approach, employing mouse Olfactory Receptors (mORs) from AlphaFold v2 as templates for homology modeling of the selected human ORs (hORs). This was done because of the higher confidence scores associated with mOR models in Alphafold, attributed to their large and validated experimental data. For OR51E2, we selected mouse Olfr78 (93% homology), Olfr558 for OR51E1 (94% homology), Olfr557 for OR51D1 (89% homology), and Olfr577 for OR51G2 (91% homology), reflecting a strategy to capitalize on the closest orthologs between mORs and hORs for enhanced model reliability.

We used MODELLER^29^ to perform this homology-based structure prediction on Alphafoldv2 models. We then compared all three models to OR51E2 and its experimental structure to ascertain the best-predicted structure obtained. We compared these models on three protein stability parameters: RMSD, MolProbity,^30^ and Ramachandran Plots. We used PyRama^31^ to assess Ramachandran plot deviations across the entire sequences, and Glycine and Proline residues as well . We used Chimera^32^ to calculate the RMSD across all protein residues and pruned pairs.

### Binding Site Prediction

In our study, we utilized PUResNet, as introduced by Kandel et al.(2021),^33^ alongside COACH, which aggregates various binding site prediction tools such as TM-SITE and ALIGN, to predict ligand-binding sites—a critical task for understanding protein interactions and functions.^34,35^ PUResNet distinguishes itself by employing structural similarity for the prediction of protein-ligand binding sites and is trained on the scPDB database, known for its annotated proteins with confirmed druggable binding sites.^36^ In the case of GPCRs, and specifically OR51E2, our comparative analysis, shown in [Fig 3(a)] revealed that PURes-Net consistently outperformed the integrated tools within COACH by reliably identifying orthosteric ligand-binding sites. This consistency in PUResNet’s predictions underscores its effectiveness and the potential advantages of leveraging structural similarities in binding site prediction (More information in Results section).

### Molecular Dynamic Simulations

Molecular Dynamic Simulations were conducted on both OR51G2 models (AFv2 and Hybrid stricture) to investigate protein-membrane interaction mechanisms and determine the more stable model for computational analysis.

The model preparation wad done using CHARMM-GUI, and the protein structures were embedded in a POPC lipid bilayer. The parameters for the protein, lipids, and ions were established using the CHARMM36m force field, and the water was modeled using the TIP3P approach. More information provided in the Supporting Information.

## Results and Discussion

The binding affinity for one of the selected potential agonists to OR51G2 was found to be the highest among all candidates. To provide a comprehensive analysis, we compared the binding behaviors of all four receptors (hOR51E1, hOR51E2, hOR51D1, and hOR51G2), offering a broad perspective on the results. We also assessed the structural stability of the OR51G2 protein models using Molecular Dyanmic Simulations. Both AlphaFold-generated models and hybrid models, which used AlphaFold structures as templates in homology modeling, underwent a detailed protein stability evaluation. This assessment encompassed multiple analytical methods, including Molecular Dynamics Simulations, to ensure a thorough examination of protein stability across different modeling approaches.

### Structure Prediction

In our study, BLAST analysis revealed sequence homologies of 59.8% for OR51E1, 58.44% for OR51D1, and 57.38% for OR51G2, indicating a considerable level of evolutionary conservation and suggesting potential functional or structural similarities among these proteins. Comparative analysis of structural models revealed the Hybrid model as the closest representation of the experimental structure, with a Root Mean Square Deviation (RMSD) of 2.238^°^A overall and 1.019^°^A across pruned pairs, outperforming the Alphafoldv2 model (RMSD: 2.499^°^A) and a generic Homology Modeling approach (RMSD: 3.885^°^A) as observed in Fig.2.

**Figure 2:**
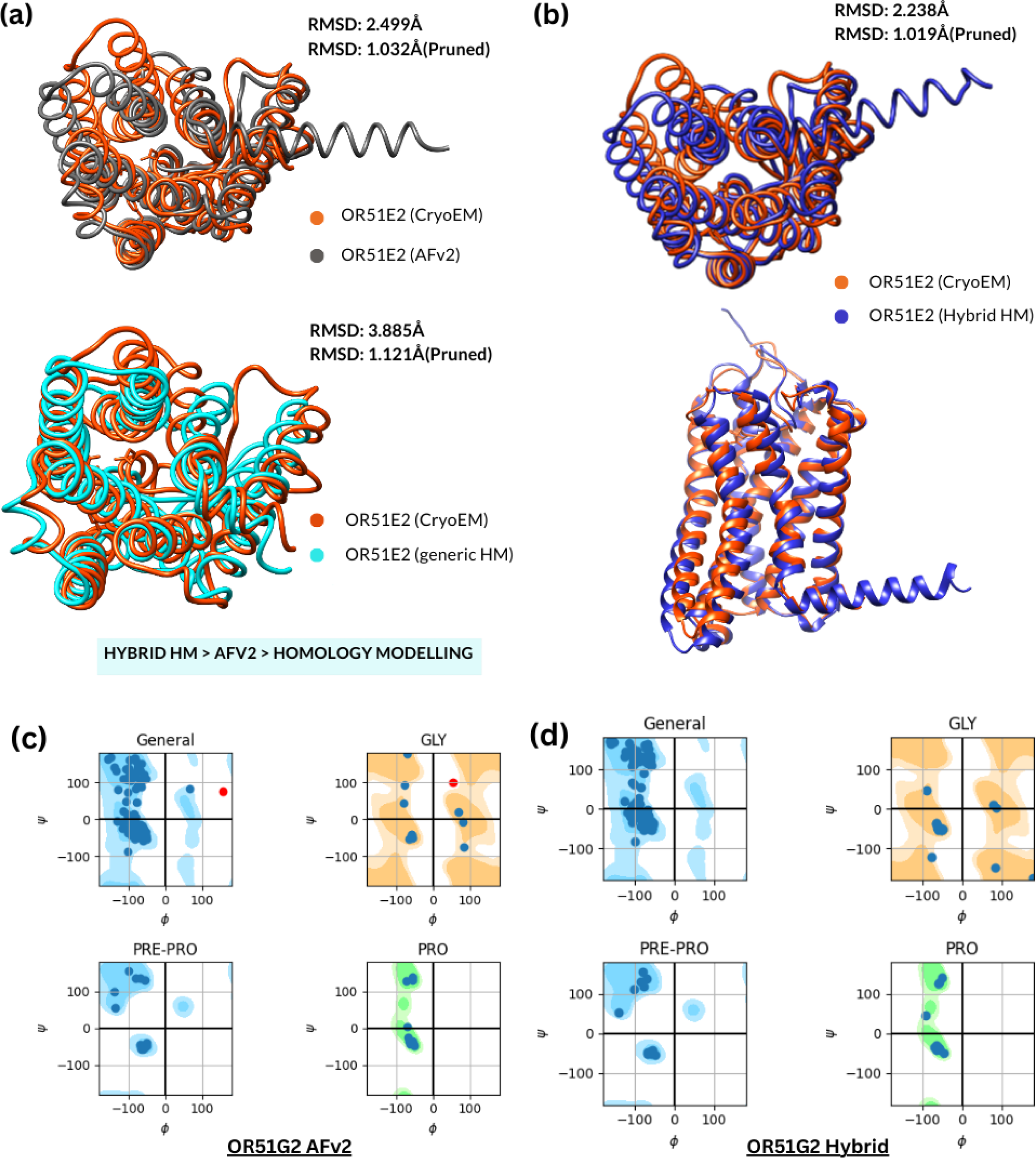
a) Shows RMSD values for AlpfafoldV2 and generic Homology Modeling performed using SwissModel and the Cryo-EM structure of an activated Cholecystokinin A receptor (CCKAR)-Gi complex. To produce an unbiased result via HM, the experimental PDB structure of OR51E2 (PDB:8F76) was not chosen as a template. (b) Showing the Hybrid Homology Modeling structure for OR51E2, modeled using Olfr78 structure from AFv2. Its high model confidence makes it a good template for modeling hORs with a very low RMSD. (c) Ramachandran plots for the two protein models in focus, OR51G2. AlphaFold models show outliers(in red) while the hybrid model has it’s residue data within the constraints.

MolProbity was chosen as the second Assessment for protein stability. We can rank the receptors based on stability and model correctness, we used the MolProbity score as the primary indicator since it is a comprehensive measure of model quality. Additionally, we considered the number of bad bonds, bad angles, and CaBLAM outliers as secondary factors. The fewer the number of deviations in these categories, the more stable and correct the model is likely to be. As observed in Table1, MolProbity score was the lowest for the Hybrid Model of OR51E2(0.62). Except in the case of OR51E1, where the Alphafold model shows a lower MolProbity score, hybrid models show better stability concerning the Alphafold structures. C*β* deviations in all models are within the thresholds, with no bad bonds and very few bad angles. Our analysis ranked the GPCR models, with the Hybrid HM OR51E2 being the most accurate, followed by Alphafold’s OR51E1, and the Experimental OR51E2 as the least.

Ranking from highest to lowest model quality:

> Hybrid HM OR51E2 *>* Alphafold OR51E1 *>* Hybrid HM OR51G2 *>* Hybrid HM OR51E1 *>* Hybrid HM OR51D1 *>* Alphafold OR51D1 *>* Alphafold OR51G2 *>* Alphafold OR51E2 *>* Experimental OR51E2.

An important observation in [Table 1] is that the Hybrid Homology Models (Hybrid HM) exhibit higher MolProbity scores compared to their AlphaFold counterparts for some of the Olfactory Receptors (ORs), yet they also show a greater number of bad angles. This seemingly paradoxical observation can be interpreted as follows:

- Higher MolProbity Scores: The higher scores in Hybrid HMs suggest that, on average, these models have better overall stereochemistry and fewer steric clashes than the AlphaFold models. This could be due to the hybrid approach incorporating additional information or constraints from experimental data or related structures during the homology modeling process, leading to improved global structural integrity.
- Increased Bad Angles: Despite the higher overall MolProbity scores, the increased number of bad angles in Hybrid HMs indicate localized deviations with bond angle geometry. This could stem from the hybrid modeling process, where the combination of template structures and computational modeling might not perfectly reconcile bond angle geometries everywhere in the structure, particularly in regions where the template and target sequences diverge significantly.^29,37^

**Table 1:** Showing the MolProbity data obtained for eight models generated from AlphaFold and the Hybrid Homology Modeling method that used high-confidence Alphafold mouse ORs as templates. It can be observed that for two out of three receptors, the Hybrid Homology Modeling method has generated a better MolProbity score. The score encompasses Clashscore, Rotamer, and Ramachandran evaluations in a single score, which then, is normalized to be on the same scale as X-ray resolution

| ORs | Model | MolProbity | Cβ dev.<br>>0.25Å | Bad<br>bonds | Bad an-<br>gles | CaBLAM |
| --- | --- | --- | --- | --- | --- | --- |
| OR51E2 | Experimental | 1.82 | 0 | 0 / 2557 | 0 / 3302 | 2 (0.7%) |
|  | AlphaFold | 1.15 | 0 | 0 / 2557 | 6 / 3478<br>(0.17%) | 1 (0.3%) |
|  | Hybrid HM | 0.64 | 1 | 0 / 2557 | 21 / 3478<br>(0.60%) | 1 (0.3%) |
| OR51E1 | AlphaFold | 0.96 | 1<br>(0.33%) | 0 / 2539 | 0 / 2539<br>(0.23%) | 0 (0.0%) |
|  | Hybrid HM | 1.10 | 1<br>(0.33%) | 0 / 2533 | 21 / 3450<br>(0.61%) | 2 (0.6%) |
| OR51D1 | AlphaFold | 1.42 | 1<br>(0.33%) | 0 / 2580 | 5 / 3514<br>(0.14%) | 1 (0.3%) |
|  | Hybrid HM | 1.23 | 0<br>(0.00%) | 0 / 2554 | 20 / 3480<br>(0.57%) | 2 (0.6%) |
| OR51G2 | AlphaFold | 1.57 | 1<br>(0.33%) | 0 / 2520 | 6 / 3423<br>(0.18%) | 2 (0.6%) |
|  | Hybrid HM | 0.99 | 0<br>(0.00%) | 0 / 2514 | 22 / 3414<br>(0.64%) | 2 (0.6%) |

Ramachandran Plot Analysis was conducted to evaluate the structural quality of the protein models. Figure 2(d) illustrates that all residues in the hybrid models fall within the favorable conformational space, indicating a high degree of structural integrity. In contrast, only two out of four of the AlphaFold models demonstrated a similar distribution of residues within the favorable range. The hybrid model of OR51E1 was an exception, presenting a few outliers; however, these were notably closer to the acceptable regions compared to the outliers observed in the AlphaFold models.

These analyses underscore greater accuracy of the Hybrid models when compared with Alphafold structures in replicating the three-dimensional structures of proteins in a manner that closely mirrors their natural biological counterparts. The observed sequence homology within the OR51-family of proteins implies their related but unique biological functions, suggesting a propensity for interacting with similar receptors. This comparative study of protein structure quality reinforces the efficacy of Hybrid modeling techniques in generating precise protein conformations.

### Binding Site Prediction

PuResNET was selected for binding site prediction after a comparison with COACH, which consistently identified the orthosteric ligand-binding sites across the olfactory receptors (ORs) under study.

The predicted binding pockets were located within the orthosteric sites, making them the focal point for subsequent analyses. PuResNET’s predictions were particularly robust for OR51E2, with all potential interacting residues aligning well within the transmembrane domains TM3, TM4, TM5, TM6, and TM7 for OR51D1 and OR51G2, while TM2 was consistently not involved in binding interactions, as represented in Figure 3(c).

**Figure 3:**
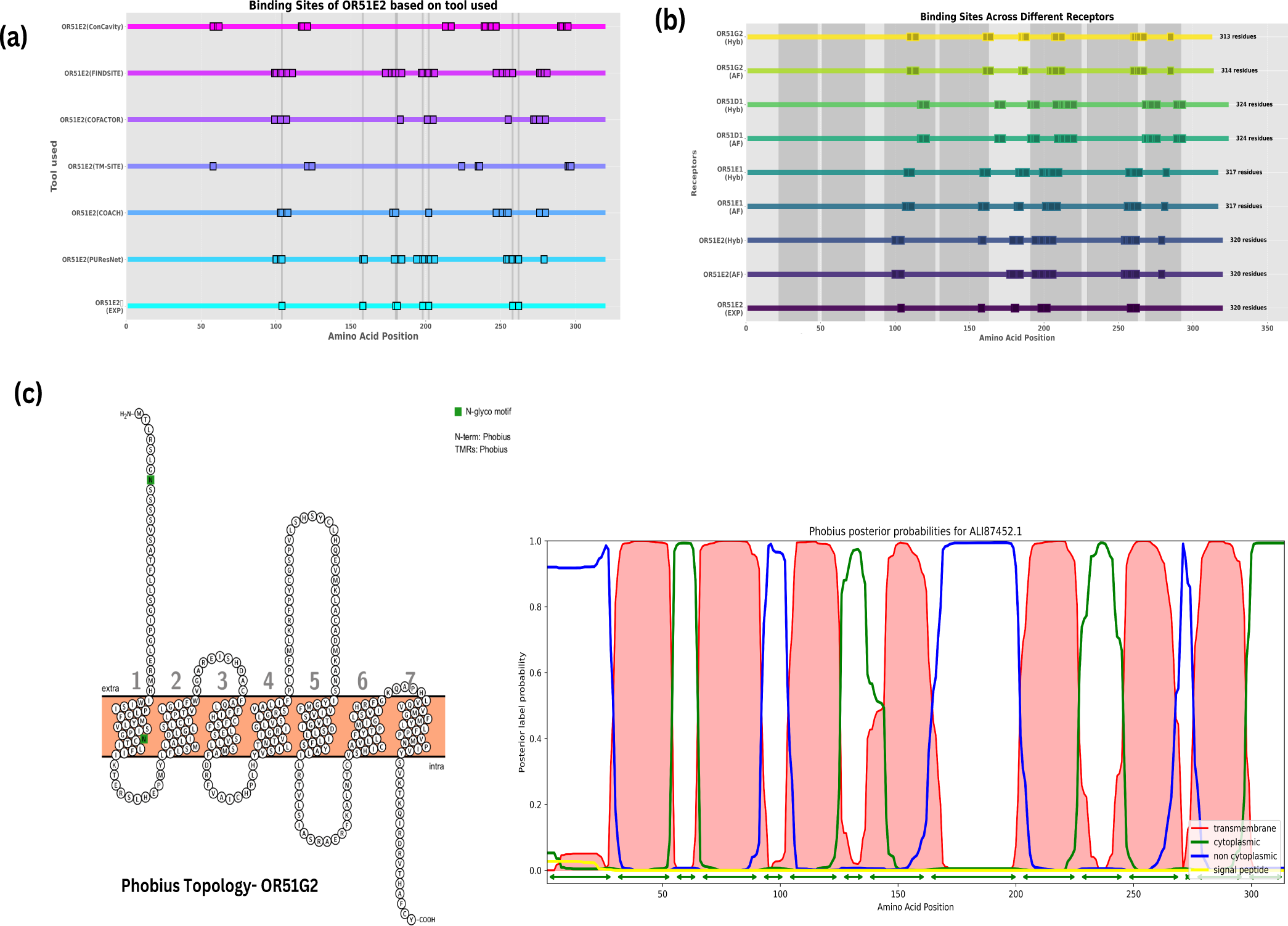
(a) The first graph shows the binding Sites of all receptors being studied across their residues. Each shaded area from left to right represents TM1-TM7, sequences based on OR51E2. It is observable that PuResNET has predicted potential binding affinity in TM7 as well. TM3-7 are the occupied ones. (b) This graph represents the comparison of the binding site predictions of OR51E2, using various methods like COACH, LIGSITE, T-SITE, etc, and compared with the experimental binding sites.(c) Topological information about OR51G2 shows what residue persists within the membrane. The Phobius Topology graph shows the same with cytoplasmic information as well.

### Molecular Dynamic Simulations

Molecular Dynamic Simulations were conducted on both OR51G2 models to investigate protein-membrane interaction mechanisms and determine the more stable model for computational analysis.

### Trajectory Analysis for OR51G2

Gromacs was used to study the stability of both models after mdrun, and the RMSD across the alpha-C backbone of the GPCRs was measured using gmx rmsd throughout runs respectively. Three runs for each OR model were performed to optimally study the OR activity across a wider sample size. Root Mean Square Deviation (RMSD) is a critical metric used to understand the extent of deviation in molecular structures such as proteins, ligands, or their complexes from a reference structure.^38^

The calculated Root Mean Squared Deviation (RMSD) in 4( graphs represent the conformational stability of the olfactory receptor OR51G2 within a palmitoyloleoylphosphatidylcholine (POPC) lipid bilayer, as predicted by two computational models: AlphaFold and a Hybrid Homology model. Essentially, RMSD measures how much a group of atoms has moved from its original position. High RMSD values typically indicate significant structural instability. This instability is manifested as conformational changes in the molecule being studied. For example, in the context of protein-membrane complexes, a high RMSD value for a protein suggests that the membrane is unstable throughout simulations.^38,39^ Below are dissections of the results of the reach run concerning RMSD.

- Run 1 Analysis: The AlphaFold prediction [red graph in Fig 4(a)] demonstrates a rapid initial increase in RMSD within the first 10,000 picoseconds (ps), suggesting an abrupt departure from the reference state, which may imply an initial relaxation or fitting of the receptor within the lipid bilayer. This is followed by a period of fluctuation, with the RMSD plateauing around 0.3 nm. The fluctuations are indicative of a dynamic equilibrium state where the protein may be sampling various conformational states. The transient peak at approximately 60,000 ps could be reflective of a temporary destabilization, such as a conformational shift or interaction with the lipid environment. Whereas, the Hybrid model [blue graph in Fig 4((a)] exhibits a more gradual increase, indicating a less abrupt conformational change initially. The overall lower RMSD suggests a closer fit to the reference structure, possibly due to the hybrid model incorporating residue co-ordinates from two different templates. The RMSD settles into a range slightly below that of the AlphaFold model, with fewer and smaller fluctuations, implying a more stable conformation throughout the simulation.
- Run 2 Analysis: The AlphaFold structure shows a lower initial RMSD, suggesting a good initial fit. However, the RMSD soon increases, surpassing the Hybrid model at around 20,000 ps. The higher RMSD and greater fluctuations may indicate a more flexible model that samples a wider conformational space, which might be critical for the receptor’s functional dynamics. While the hybrid model remains relatively stable with a generally lower RMSD throughout the run. A pronounced spike in the RMSD is observed at around 80,000 ps, but it quickly returns to the previous level. This could represent a specific event, such as a conformational transition or an interaction with a lipid molecule that is resolved quickly, returning the protein to its previous state of stability.
- Run 3 Analysis: Initially, the AlphaFold model’s RMSD is slightly higher than that of the Hybrid model, but as the simulation progresses, the AlphaFold RMSD slightly decreases and remains below the Hybrid’s RMSD for most of the simulation time. The lower RMSD suggests that, in this run, the AlphaFold model is sampling conformations closer to the reference. The fluctuations are pronounced and consistent, indicating ongoing dynamic behavior and adaptation to the lipid bilayer environment. In the case for the hybrid structure, the initial RMSD is low, but it soon surpasses the AlphaFold’s RMSD and remains higher for the majority of the simulation. The consistently higher RMSD could suggest that the Hybrid model, in this specific run, is less stable or it could be more dynamic within the lipid bilayer.

**Figure 4:**
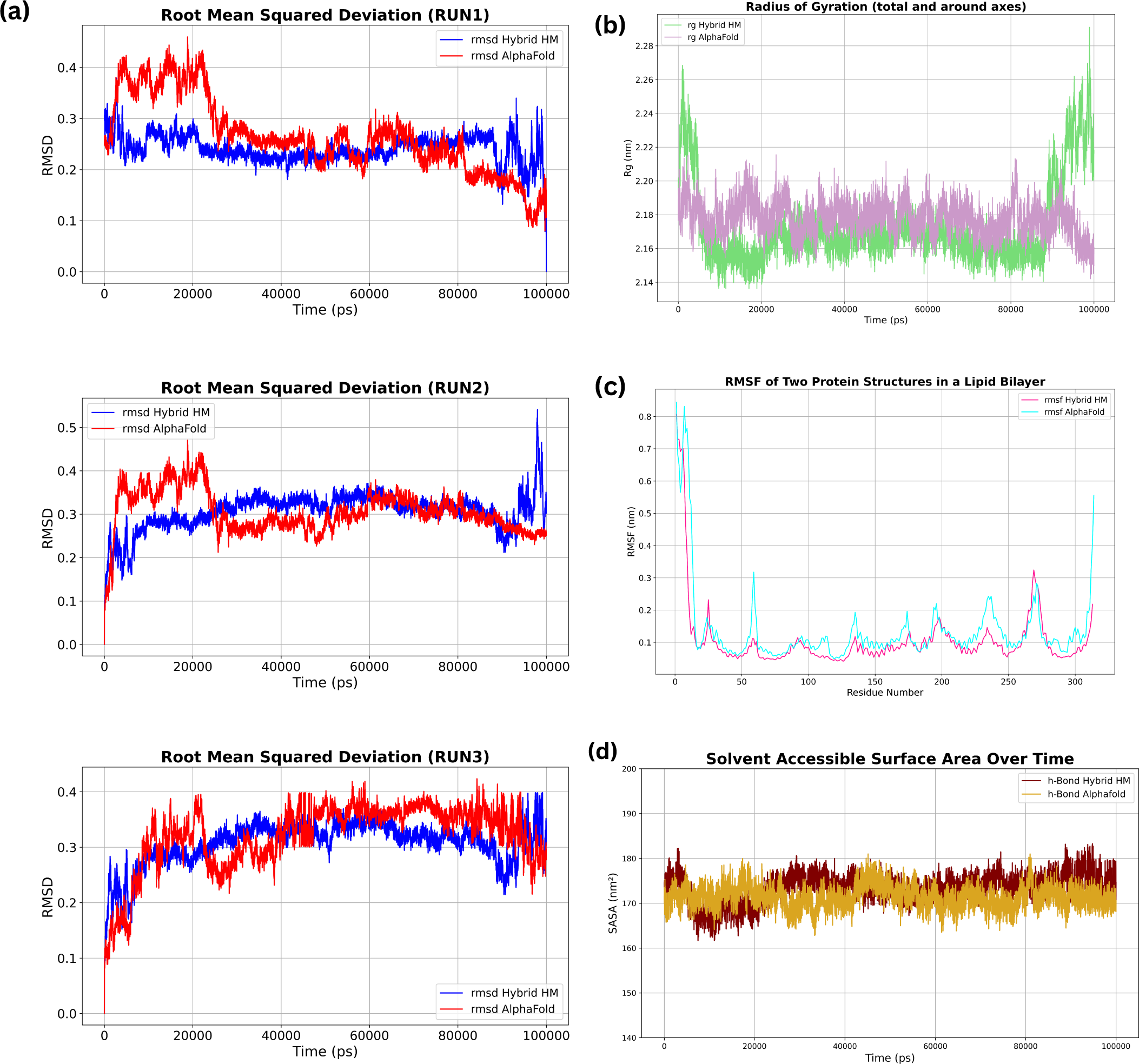
(a) Each graph shows the RMSD for three runs each for both models in comparison to each other across a length of 100 ns simulations. The Hybrid model performs better than Alphafold model in 2 runs out of three. The system for Alphafold is observed to be more flexibile initially in the first 30,000 ps as compared to an overall stable Hybrid model. (b) Radius of Gyration across the axes of the models show that hybrid model tends to have more movement in the beginning and end of the simulation. (c) the Root Mean squared Fluctuations across the receptor sequence within the lipid bilayer shows more movements aross the ECL regions present outside the memrbane in the Hybrid model exhibiting a more robust GPCR dynamics which men that the system for Hybrid OR is active and is interacting well with the extra-membranous environment. (d) the SASA calculations across the length of the simulation signify that the hybrid model is slightly more exposed to the solvent in the environment *i.e.* water.

Throughout the three simulation runs, the AlphaFold-predicted model generally exhibits higher RMSD values when compared to the Hybrid Homology model. This observation suggests that the AlphaFold model tends to deviate more from the initial conformation, reflecting either an intrinsic flexibility within the receptor or a dynamic adaptation to the membrane environment.

Specifically, the pronounced peaks and valleys in the RMSD profile of the AlphaFold model indicate that the receptor may be sampling a diverse conformational landscape. This could be a manifestation of the inherent flexibility required for the receptor to interact with diverse ligands or to undergo conformational changes that are essential for signal transduction processes. The temporal RMSD increases, particularly noted in Run 1 around 60,000 ps and in Run 2 between 40,000 to 60,000 ps, may correspond to significant structural rearrangements or interactions with lipid molecules that could be critical for the functional activity of the receptor.

In contrast, the Hybrid Homology model maintains a lower and more consistent RMSD in most instances, suggesting a conformation that is more restrained or a model that is more reflective of a stable state within the lipid bilayer. This indicates that AFv2 predicted proteins with ample experimental data, combined with experimental structures may produce predict a receptor conformation that is closer to a functional state as generally observed in experimental conditions, shown by the Hybrid structures.

In the third run, the AlphaFold model demonstrates a lower RMSD compared to the Hybrid model, inverting the trend observed in the first two runs. This underscores the stochastic nature of protein dynamics and the influence of initial conditions and random thermal fluctuations on simulation outcomes.

The diverse RMSD behaviors between the two models also raise intriguing questions regarding the accuracy and predictive power of computational models. While the AlphaFold model may capture a broader conformational space due to its purely predictive nature, the Hybrid Homology model, may provide a more restrained prediction that could either indicate a more stable receptor conformation or overlook relevant dynamic states.^40^

Radius of gyration (Rg) measurements provide further clarity on the compactness or expansion of the receptor structure over time. Solvent-accessible surface area (SASA) analysis reflects the receptor’s hydrophobic and hydrophilic surface exposure, informing on the receptor-lipid interactions below:

- Radius of Gyration (Rg) Analysis: The Rg metric serves as an indicator of the protein’s overall compactness and its structural integrity over time.^41,42^ In the graph provided, the Rg values for the AlphaFold model remain relatively stable, with minor fluctuations throughout the simulation, suggesting a consistent tertiary structure. This stability implies that the AlphaFold-predicted structures possess resilience against conformational changes within the lipid bilayer environment, essential for maintaining functional activity. Additionaly, the Rg values for the Hybrid Homology model exhibit greater variability, with significant peaks, especially towards the latter part of the simulation. The peaks observed in Fig.4(b) reflect transient unfolding events or major conformational rearrangements that could impact the protein’s functional state. Such behavior might be indicate receptor undergoing adaptive changes, possibly in response to interactions with the lipid bilayer or internal dynamics that may be essential for receptor activation or signaling.^43^
- Root Mean Square Fluctuation (RMSF) Analysis: The RMSF graph in Fig.4(c) illustrates the flexibility of each residue throughout the simulation. Both models present variable fluctuations along the residue sequence. Peaks in RMSF are commonly associated with loop regions or flexible domains that may play a crucial role in the receptor’s functionality, such as ligand-binding sites or regions important for protein-protein interactions. The AlphaFold model demonstrates certain residues with notably higher fluctuations compared to the Hybrid model, suggesting these regions have increased mobility. This increased flexibility could indicate a dynamic nature of these regions, potentially facilitating the receptor’s interactions with ligands or other cellular components.^44^ The Hybrid Homology model’s generally lower RMSF values suggest a constrained prediction of residue movements, which may imply a model that emphasizes the stability of the receptor’s structure over its dynamic properties.
- Solvent Accessible Surface Area (SASA) Analysis: SASA is a direct measure of the protein’s interaction with the solvent environment and indicates potential binding and active sites. The graph in Fig4(d) shows the SASA for both models fluctuating for the simulation. The AlphaFold model tends to display higher SASA values, which could suggest a structure with more exposed hydrophilic surfaces that may interact with solvent molecules. A consistently higher SASA might also imply a conformation with more flexible or disordered regions, potentially related to the receptor’s functional states.^45,46^

The consistent Rg and elevated SASA values in the AlphaFold model suggest a protein that maintains its structural core while presenting a surface that is more exposed and possibly more interactive with the surrounding lipid bilayer. This could be indicative of a functionally active receptor conformation, with the necessary flexibility to allow for ligand binding and subsequent conformational changes essential for signal transduction.

The Hybrid Homology model’s variable Rg and lower SASA values suggest a protein with more dynamic structural integrity but with less overall exposure to the solvent. This might reflect a receptor state that is less active or in a different functional state, possibly a resting or inactive conformation. The lower flexibility in certain regions as indicated by RMSF might also suggest a receptor that is more rigid in its interactions within the cellular environment.

The biophysical characteristics depicted by these models offer an insight on the quality and stabilities of both these models. The Alphafold model shows inconsistent deviations across simulations and high fluctuations along the length of the protein, suggesting deviations from standard GPCR behavior, whereas the hybrid model shows consistent behavior as compared to Alphafold, as it has more stable RMSD across the simulations.

### Conclusion

In this study, we developed a comprehensive Computer-Assisted pipeline to investigate olfactory receptor (OR)-odorant interactions, with a focus on the OR51-family. Our approach integrated advanced computational techniques, including hybrid homology modeling, and molecular dynamics simulations, to predict the structure of specific ORs. Our findings demonstrate that the hybrid homology modeling strategy, which leveraged Olfrs from *Mus musculus* in this case, produced more accurate and stable models of ORs compared to AlphaFold predictions alone. We obtained significant improvements in structural metrics such as RMSD, MolProbity scores, and Ramachandran plot analysis. The enhanced accuracy of these models was crucial for reliable prediction of OR-odorant interactions, as demonstrated by our successful replication of experimental findings for OR51E2. The hybrid models show consistent RMSD values, stable radius of gyration, and realistic flexibility in key regions, indicating their suitability for further computational studies. This workflow can be potentially expanded over to other GPCRs as well.

## Data Availability Statement

The raw data supporting the conclusions of this article will be made available by the authors, without undue reservation.

## Supplementary Material

The Supplementary Material for this article can be found online as repository: Structure Analysis Data

## Acknowledgements

The authors thank CSIR for funding the research.

## References

(1) Sanne, B.; Valentina, P. The importance of the olfactory system in human well-being, through nutrition and social behavior. Cell Tissue Res. 2021, 383, 559–567.

(2) Jinyoung, S.; Subin, C.; Hyeyoun, K.; See-Hyoung, P.; Jongsung, L. Association between Olfactory Receptors and Skin Physiology. Journal of the American Chemical Society 2022, 135, 7296–7303.

(3) Maßberg, D.; Hatt, H. Human Olfactory Receptors: Novel Cellular Functions Outside of the Nose. Physiological reviews 2018, 98 *3*, 1739–1763.

(4) Sarafoleanu, C. The importance of the olfactory sense in the human behavior and evolution. BMB reports 2009, 2.

(5) Kang, N.; Koo, J. Olfactory receptors in non-chemosensory tissues. BMB reports 2012, 45.

(6) Nielsen, B. L. Olfaction: An Overlooked Sensory Modality in Applied Ethology and Animal Welfare. BMB reports 2015, 2.

(7) Yoshihito, N.; Atsushi, M.; Kazushige, T. Recent advances in the determination of G protein-coupled receptor structures. Cell 2014, 24.

(8) Obot, D. N.; J. Udom, G. e. Advances in the molecular understanding of G proteincoupled receptors and their future therapeutic opportunities. Futur J Pharm Sci 2021, 7.

(9) Orecchioni, M. e. Olfactory receptor 2 in vascular macrophages drives atherosclerosis by NLRP3-dependent IL-1 production. Science 2022, 375.

(10) Almeida, J. G.; Preto, A. J.; Koukos, P. I.; Bonvin, A. M.; Moreira, I. S. Membrane proteins structures: A review on computational modeling tools. Biochimica et Biophysica Acta (BBA) - Biomembranes 2017, 1859, 2021–2039.

(11) Alberts, B.; Johnson, A.; Lewis, J.; Raff, M.; Roberts, K.; Walter, P. Molecular Biology of the Cell, 4th ed.; Garland Science: New York, 2002; Chapter Membrane Proteins.

(12) Shimada, I.; Ueda, T.; Kofuku, Y.; Eddy, M. T.; Wüthrich, K. GPCR drug discovery: integrating solution NMR data with crystal and cryo-EM structures. Nature Reviews Drug Discovery 2018, 18, 59–82.

(13) Latorraca, N. R.; Venkatakrishnan, A. J.; Dror, R. O. GPCR Dynamics: Structures in Motion. Chemical Reviews 2017, 117, 139–155.

(14) Esguerra, M.; Siretskiy, A.; Bello, X.; Sallander, J.; Gutiérrez-de Terán, H. GPCR-ModSim: A comprehensive web based solution for modeling G-protein coupled receptors. Nucleic Acids Research 2016, 44, W455–W462.

(15) Ranganathan, A.; Dror, R. O.; Carlsson, J. Insights into the Role of Asp79(2.50) in 2 Adrenergic Receptor Activation from Molecular Dynamics Simulations. Biochemistry 2014, 53, 7283–7296.

(16) Jumper, E. R. P. A. e. J. Highly accurate protein structure prediction with AlphaFold. Nature 2021, 596.

(17) Sala, D.; Hildebrand, P. W.; Meiler, J. Biasing AlphaFold2 to predict GPCRs and kinases with user-defined functional or structural properties. Frontiers in Molecular Biosciences 2023, 10, 1121962.

(18) Cong, X.; Ren, W.; Pacalon, J.; Xu, R.; Xu, L.; Li, X.; de March, C. A.; Matsunami, H.; Yu, H.; Yu, Y.; Golebiowski, J. Large-Scale G Protein-Coupled Olfactory Receptor–Ligand Pairing. ACS Central Science 2022, 8, 379–387.

(19) Spehr, M.; Munger, S. D. Olfactory receptors: G protein-coupled receptors and beyond. Journal of Neurochemistry 2009, 109, 1570–1583.

(20) Slabinski, L.; Jaroszewski, L.; Rodrigues, A. P.; Rychlewski, L.; Wilson, I. A.; Lesley, S. A.; Godzik, A. The challenge of protein structure determination–lessons from structural genomics. Protein Science 2007, 16, 2472–2482.

(21) Hassell, A. M. et al. Crystallization of protein-ligand complexes. Acta Crystallographica Section D: Biological Crystallography 2007, 63, 72–79.

(22) Leki, T.; Yamanak, T.; Yoshikawa, K. Functional analysis of human olfactory receptors with a high basal activity using LNCaP cell line. PLoS One 2022, 17.

(23) TAariq, M. U. e. a. Methods for Proteogenomics Data Analysis, Challenges, and Scalability Bottlenecks: A Survey. IEEE 2020, 9.

(24) Hess, J.; Kohl, T.; Kotrová, M.; Rönsch, K.; Paprotka, T.; Mohr, V.; Hutzenlaub, T.; Brüggemann, M.; Zengerle, R.; Niemann, S.; Paust, N. Library preparation for next generation sequencing: A review of automation strategies. Biotechnology Advances 2020, 41, 107537.

(25) Alfonso-Prieto, M.; Capelli, R. Machine Learning-Based Modeling of Olfactory Receptors in Their Inactive State: Human OR51E2 as a Case Study. J. Chem. Inf. Model. 2023, 63.

(26) Billesbølle, C. B.; March, C. A. d.; van der Velden, W. J. C. e. Structural basis of odorant recognition by a human odorant receptor. Nature 2023, 615.

(27) Liu, X. e. The number of protein folds and their distribution over families in nature. Proteins *vol.* 2004, 54.

(28) Waterhouse, A. e. SWISS-MODEL: homology modelling of protein structures and complexes. Nucleic Acids Research 2018, 46.

(29) Webb, B.; Sali, A. Comparative Protein Structure Modeling Using MODELLER. Current protocols in bioinformatics 2016, 54.

(30) Williams, C. J. e. a. MolProbity: More and better reference data for improved all-atom structure validation. Proteins Science - Wiley 2018, 27.

(31) Lovell, S. C. e. Structure validation by C geometry:, and C deviation. Proteins – Wiley 2003, 50.

(32) Pettersen, E. e. UCSF Chimera–a visualization system for exploratory research and analysis. J Comput Chem. 2004, 25.

(33) Kandel, J.; Tayara, H.; Chong, K. T. PUResNet: prediction of protein-ligand binding sites using deep residual neural network. Journal of Cheminformatics 2021,

(34) Yang, J. e. a. Protein–ligand binding site recognition using complementary binding-specific substructure comparison and sequence profile alignment. Bioinformatics - Oxford 2013, 29.

(35) Yang, J.; Roy, A.; Zhang, Y. BioLiP: a semi-manually curated database for biologically relevant ligand–protein interactions. Nucleic Acids Research 2012, 41, D1096–D1103.

(36) Jérémy, D. e. sc-PDB: a 3D-database of ligandable binding sites—10 years on. Nucleic Acids Res. 2015, 43.

(37) Davis, I. W. e. a. MolProbity: all-atom contacts and structure validation for proteins and nucleic acids. Nucleic Acids Research-Oxford 2007, 35.

(38) Hospital, A. e. Molecular dynamics simulations: advances and applications. Frontiers in physiology 2015, 8, 37–47.

(39) Sargsyan, K. e. How Molecular Size Impacts RMSD Applications in Molecular Dynamics Simulations. ACS 2017, 13, 37–47.

(40) Lindahl, E.; Sansom, M. S. Membrane proteins: molecular dynamics simulations. Current Opinion in Structural Biology 2008, 18, 425–431, Membranes / Engineering and design.

(41) Lobanov, M. e. Radius of gyration as an indicator of protein structure compactness. Molecular Biology 2008, 42, 623–628.

(42) Arnittali, M. e. Structure Of Biomolecules Through Molecular Dynamics Simulations. Procedia Computer Science 2019, 156.

(43) Maity, S. e. Conformational rearrangements in the transmembrane domain of CNGA1 channels revealed by single-molecule force spectroscopy. Nature Communications 2015, 6.

(44) Malik, N. e. Strategies for identifying dynamic regions in protein complexes: Flexibility changes accompany methylation in chemotaxis receptor signaling states. Biochimica et Biophysica Acta (BBA) - Biomembranes 2020, 1862.

(45) Harroun, S.; Lauzon, D. Monitoring protein conformational changes using fluorescent nanoantennas. Nature Methods 2021, 19, 71–80.

(46) Wei, S. e. A rapid solvent accessible surface area estimator for coarse grained molecular simulations. Journal of computational chemistry 2017, 38.

